# Zoonotic *Babesia microti* in the northeastern U.S.: evidence for the expansion of a specific parasite lineage

**DOI:** 10.1101/223420

**Authors:** Heidi K. Goethert, Philip Molloy, Victor Berardi, Karen Weeks, Sam R. Telford

**Author notes:** Corresponding author (SRT).

## Abstract

The recent range expansion of human babesiosis in the northeastern United States, once found only in restricted coastal sites, is not well understood. This study sought to utilize a large number of samples to examine the population structure of the parasites on a fine scale to provide insights into the mode of emergence across the region. 228 *B. microti* samples collected in endemic northeastern U.S. sites were genotyped using published VNTR markers. The genetic diversity and population structure were analysed on a geographic scale using Phyloviz and TESS. Three distinct populations were detected in northeastern US, each dominated by a single ancestral type. In contrast to the limited range of the Nantucket and Cape Cod populations, the mainland population dominated from New Jersey eastward to Boston. Ancestral populations of *B. microti* were sufficiently isolated to differentiate into distinct populations. Despite this, a single population was detected across a large geographic area of the northeast that historically had at least 3 distinct foci of transmission, central New Jersey, Long Island and southeastern Connecticut. We conclude that a single *B. microti* genotype has expanded across the northeastern U.S. The biological attributes associated with this parasite genotype that have contributed to such a selective sweep remain to be identified.

**Author summary:** Babesiosis is a disease caused by a protozoan parasite, *Babesia microti,* related to malaria. The disease is acquired by the bite of the deer tick, the same tick that transmits Lyme disease. Although Lyme disease rapidly emerged over a wide range within the last 40 years, babesiosis remained rare with an extremely focal distribution. Within the last decade, the number of reports of babesiosis cases has increased from an expanded area of risk, particularly across the mainland of southern New England. We determined whether the expanded risk may be due to local intensification of transmission as opposed to introduction of the parasite. Historical fragmentation of the landscape suggests that sites of *B. microti* transmission should have been isolated and thus evidence of multiple genetically distinct populations should be found. By a genetic fingerprinting method, we found that samples from the new mainland sites were all genetically similar. We conclude that one parasite genetic lineage has recently expanded its distribution and now dominates, suggesting that it has some phenotypic attribute that may confer a selective advantage over others.

## Introduction

Human babesiosis due to *Babesia microti* was first recognized on Nantucket Island nearly 50 years ago [1], and a few years later the first cases of Lyme arthritis were described from Old Lyme, Connecticut [2]. Both infections were found to be transmitted by the deer tick *(Ixodes dammini;* American clade of *I. scapularis),* which had started to be locally recognized as a human-biting pest [3]. In the 1970s and 80s, cases of either were restricted to coastal New England sites, as well as foci in Wisconsin and Minnesota [4–6]. Over the next 20 years, the number of Lyme disease cases significantly increased and zoonotic risk spread rapidly across the northeastern United States. Lyme disease is now endemic all the way north into Canada, west to Ohio, and south as far as Virginia. Babesiosis, in constrast, lagged Lyme disease across these sites in time and in force of transmission [7, 8] and most cases were reported from coastal sites in the northeastern U.S. However, in the last two decades, risk for babesiosis has intensified across the northeastern U.S. [9, 10].

The 20 year lag between the range expansion of Lyme disease and that of babesiosis is not fully understood but in part relates to the difficulty with which *B. microti* may be transported. The two key facts that pose a paradox for range expansion are (1) only rodents and insectivores are known to be competent reservoirs of *B. microti* (may pass infection to uninfected ticks; [11]; and (2) *B. microti* is not inherited by ticks [11]. Larval ticks transported long distances by migratory birds, a critical mode of introduction for the agent of Lyme disease (for which certain passerines are competent reservoirs; [12]), do not develop into infected nymphs after they engorge on a bird because birds are not likely to be reservoir competent for *B. microti.* A *B. microti-infected* nymph (which acquired infection as a larva feeding on a mouse) transported by a bird could develop into an infected adult tick, but because that stage feeds only on medium to large sized mammals, especially deer, would not pass infection to a reservoir competent animal during the adult bloodmeal; deer are not competent reservoirs and carnivores are not likely to be competent. Hence, *B. burgdorferi* is said to travel on the backs of birds but *B. microti* on mice. Mice or other small mammals are unlikely to travel large distances. These considerations argue that the range expansion for *B. microti* babesiosis is not due to introductions of infected ticks by migratory birds.

The existence of silent natural foci of transmission is suggested by early rodent serosurveys for *B. microti* in Connecticut [13] and the detection of zoonotic clade parasites from sites in Maine where human babesiosis had not been recorded [14]. However, ecological surveillance has not been conducted across the northeastern U.S. with sufficient detail across the likely range to provide much data of utility in understanding the tempo and mode of babesiosis risk. Longitudinal analyses of cases reported to state departments of public health are useful because case reports are based on a standard surveillance case definition and data are comparable between states. In Rhode Island, risk diminished from south to north [15]. In New York, babesiosis case reports gradually expanded from Long Island up the Hudson River valley. Similarly, in Connecticut, case reports expanded through the years from the southeastern coast first extending westward along the coast and then moving inland. [7,16–18]. The expansion of risk has been limited and incremental, with no long-distance introduction events such as those documented for Lyme disease, exemplified by its introduction into Canada. [19] A recent model for the emergence of babesiosis in New England suggests a “stepping-stone” model: a strong predictor of a town reporting babesiosis cases was the presence of a neighboring town reporting cases and that Lyme disease risk was a prerequisite [8]. Two stepping stone scenarios might have been operating concurrently in the last 20 years. (1) The force of *B. microti* transmission increased, slow and wave-like, across the northeastern landscape with the coastal earliest known zooonotic sites seeding adjacent more northerly sites. (2) Multiple cryptic enzootic sites (natural foci) with little zoonotic risk existed across the region, with local intensification of the force of *B. microti* transmission as tick densities increased to a threshold (estimated to be more than 20 nymphal deer ticks collected per hour; [20], and subsequent spread to adjacent areas.

The population structure of *B. microti* may provide evidence for tempo and mode of the expansion of babesiosis risk across the northeast. At the very basic level, new demes will be related genetically to their parent populations. In expanding populations, genetic diversity may be low be due to bottlenecks and founder effects at the expanding front [21, 22]. In fact, observed patterns of diversity will vary depending on the process of population expansion, viz., whether the population is being “pushed” or “pulled”[21, 23]. A “pulled” expansion occurs when pioneers are seeding new populations ahead of the source population, such as would occur if individual infected ticks are being introduced into a new site. This causes the genetic diversity to be lower at the edge than the main body of the population due to successive founder effects. By contrast, a “pushed” expansion occurs when a population expands at the edges of the source location due to population growth. This expansion is usually slower and allows for diversity in the source population to keep pace with geographical spread. A skewed population diversity can occur near the expanding front of the population due to “allele surfing”, that is high rates of reproduction can increase mutation and allow an allele to surf the wave of population growth and become prevalent when it might not have become fixed in a stationary population [21,24–26].

We have previously described variable number tandem repeat (VNTR) markers for analyzing the population structure of *B. microti* and detected 3 distinct populations in ticks and rodents across New England [27]. Whole genome sequencing of ecological and clinical samples determined that these *B. microti* populations were strongly differentiated, suggesting that they were geographically isolated [28]. However, neither study analyzed sufficient samples to provide detail on the mode of expansion of the range of B.microti in the northeastern U.S. Accordingly, we leveraged >200 diagnostic blood samples from patients suspected of having acute babesiosis presenting to several clinical practices across the northeastern U.S. and analyzed them with the VNTR assay. In particular, we sought to determine the population structure of these parasites, and whether range expansion was best represented by a “pulled” expansion model by introductions into small founder populations, or a “pushed” model consistent with stepping stone expansion.

## Materials and methods

### *B. microti* blood samples

De-identified discarded blood samples were collected from specimens that had been sent to Imugen, Inc. for diagnosis of *B. microti* infection during the transmission season of 2015. The town of the submitting doctor’s office or hospital was associated with each sample but no other data was available. Samples with a Ct>34 on the diagnostic real time PCR performed at Imugen were excluded from the analysis because they would not have had enough parasite DNA to yield reliable VNTR typing results.

### Ethics Statement

This study was considered not to comprise human subjects research by the Tufts University institutional review board.

### Genotyping

DNA was extracted using a commercial spin column method (Qiagen Inc.). *B. microti* was typed as described [27], with the exception that the hypervariable locus, BMV4, was excluded. Samples were excluded from the final analysis if more than 1 locus failed to amplify. To avoid erroneously scoring stutter peaks, multiple peaks were scored only if the size of the minor peak was almost equal to that of the major peak. *B. microti* merozoites infecting humans are haploid [29]; so all analyses were done under the assumption of haploidy. Samples that had multiple peaks in more than one locus were excluded, as it was impossible to determine the individual haplotypes needed for assigning a haplotype to a population using Phyloviz (see below). Samples that had multiple peaks in only a single locus were retained in the analysis and treated as two separate haplotypes.

### Data analysis

VNTR haplotypes were analyzed with two programs (Phyloviz [30] and TESS [31]) that utilize different algorithms for assigning them to a population. Phyloviz determines mutually exclusive related groups by using the eburst algorithm on haplotype data to identify founder haplotypes and then predicts the descent from the founder to the other haplotypes without any predefined assumptions of populations or geographic location. TESS uses a Bayesian clustering algorithm to determine population structure from geographically defined haplotypes without assuming predefined populations.

TESS requires that a unique geographic location be associated with each sample. Because samples were de-identified and only the location of the contributing clinical practice was known, we created random locations for each sample within a standard deviation of 0.05 degrees longitude and 0.025 degrees latitude from the town associated with each sample using the tool provided by TESS. To ensure that nearest neighbor connections could not occur over the ocean, 23 dummy points, i.e. points at which sampling cannot occur, were added in the Atlantic Ocean along the shoreline. In addition, the spatial network was altered to remove any remaining nearest neighbor connections that spanned the ocean. Geographic distances between each sample point were calculated using TESS. The program was then run for 10 permutations for K populations, from 2 to 8, with allowance for admixture. The mean deviance information criterion (DIC) was calculated across runs for each K population in order to choose the best fit among alternate models. The output from the 10 individual runs of the chosen K was downloaded into CLUMPP [32] which compiled them together. The resulting ancestry coefficients were displayed as a bar graph. An ancestry coefficient of 0.80 or greater for a single population was determined to be a member of that population. Any sample with a coefficient less than 0.80 for any single population was determined to have significant admixture from more than 1 source population. The ancestry coefficients were spatially interpolated onto a map of New England using R [33]. Fst estimates were calculated with Genepop on the web [34, 35], PhiPT estimates were calculated using GenAlEx [36], and the Shannon Index of Diversity was calculated using PAST [37] on samples grouped by region. The Outline map of the northeastern United States was downloaded from dmaps.com (http://dmaps.com/carte.php?num_car=3895&lang=en)

## Results

*B. microti* was typed from 234 specimens from 24 towns throughout New England during 2015 (Fig 1 and Table 1). Of these samples, 42 had multiple alleles in one locus and 6 had multiple alleles for more than 1 locus. The latter were excluded from the analysis because we were unable to accurately determine the haplotype necessary for analysis by Phyloviz. From the 228 samples used in the study, 113 unique haplotypes were obtained. The samples were grouped by geographic region (Table 1) and the Shannon Index (H) was calculated for each region (Fig 2). The diversity for most regions ranged from 1.8-2.5 and was not significantly different from each other. However, the diversity from the New Jersey (NJ) samples was significantly lower (H=0.9, p=0.02) and the diversity from southeastern Massachusetts (SeMA) samples was significantly higher (H=3.3, p<0.001) than the rest. Population differentiation estimates, PhiPT, suggest isolation between some regions and almost none between others (Table 2). Samples from Nantucket (N) and Cape Cod (CC) have significant amounts of population differentiation between each other and each of the other geographic groups. (Table 2) In contrast, there is no evidence of any population differentiation between samples from NJ, Long Island (LI), Connecticut (CT) and Rhode Island (RI). Samples from SeMA and western Massachusetts (WMA) show moderate amounts of population differentiation between each other and those from NJ, LI, CT and RI (Table 2).

**Figure 1.**
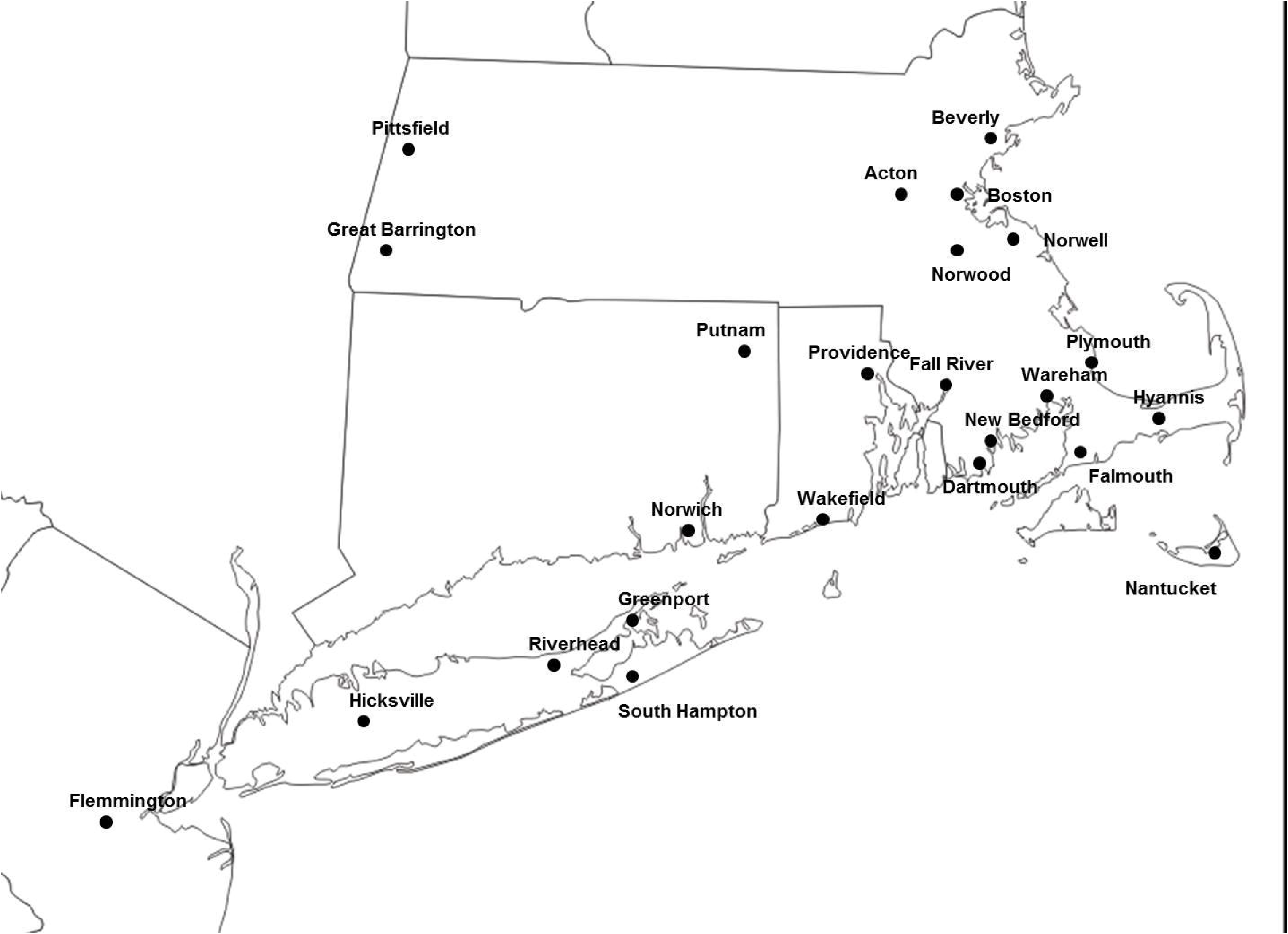
Map of the Northeastern United States labeled with the sites from which samples were collected.

**Figure 2.**
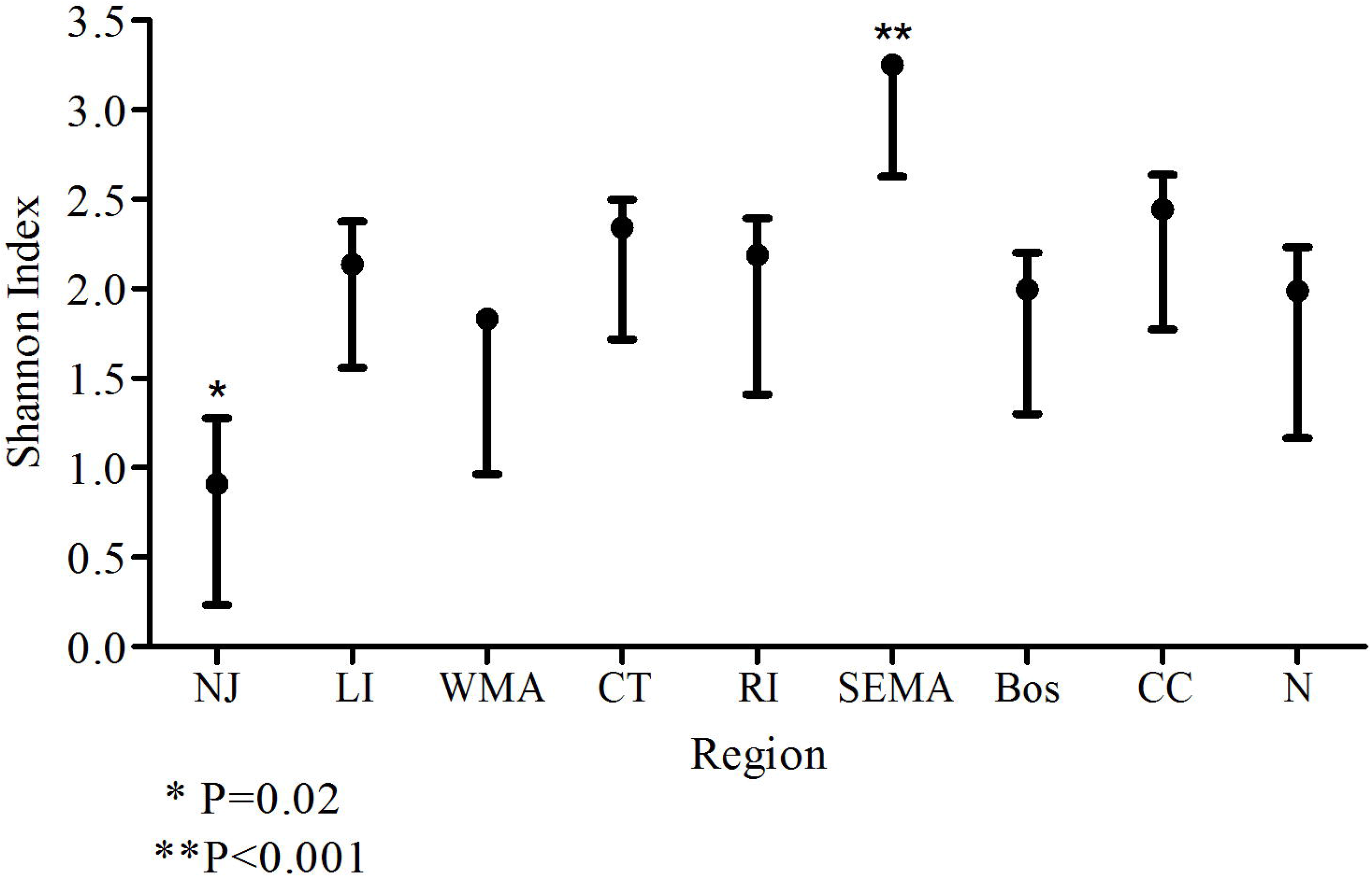
Shannon’s Index of diversity with standard error for *B. microti* haplotypes found in each region. New Jersey (NJ), Long Island (LI), Western Massachusetts (WMA), Connecticut (CT), Rhode Island (RI), southeastern Massachusetts (SEMA), Boston (Bos), Cape Cod (CC) and Nantucket (N).

**Table 1.**
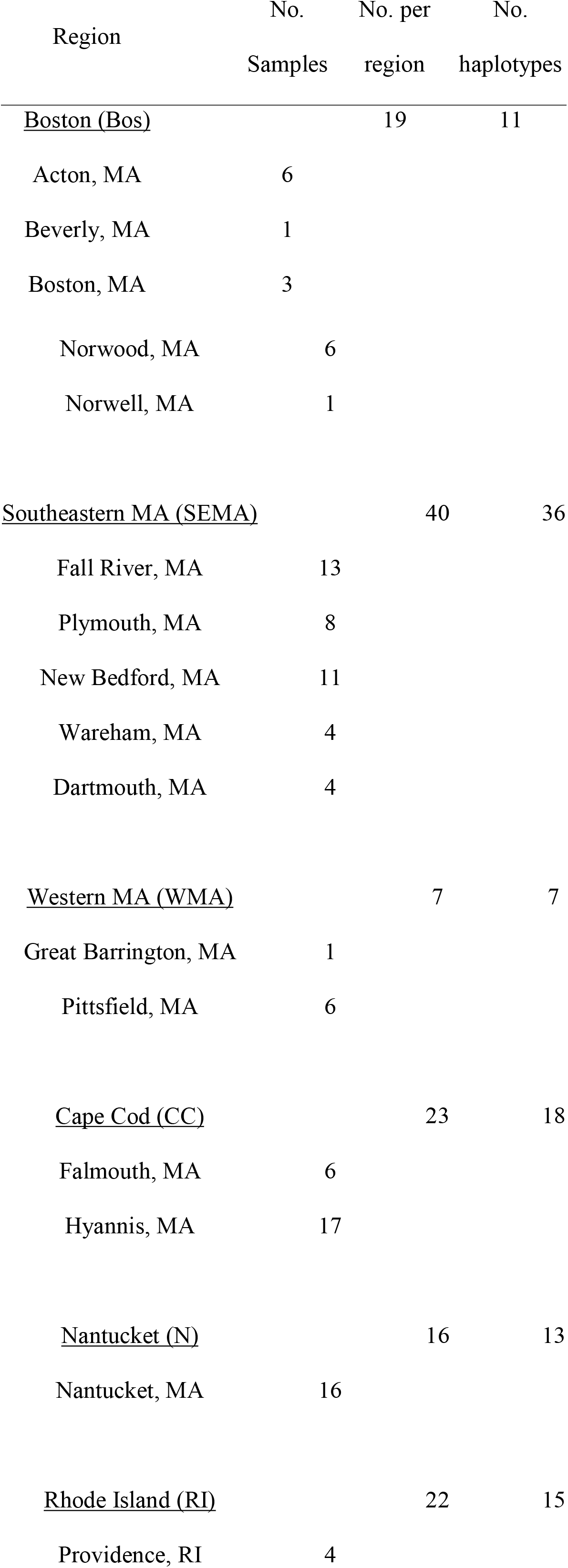
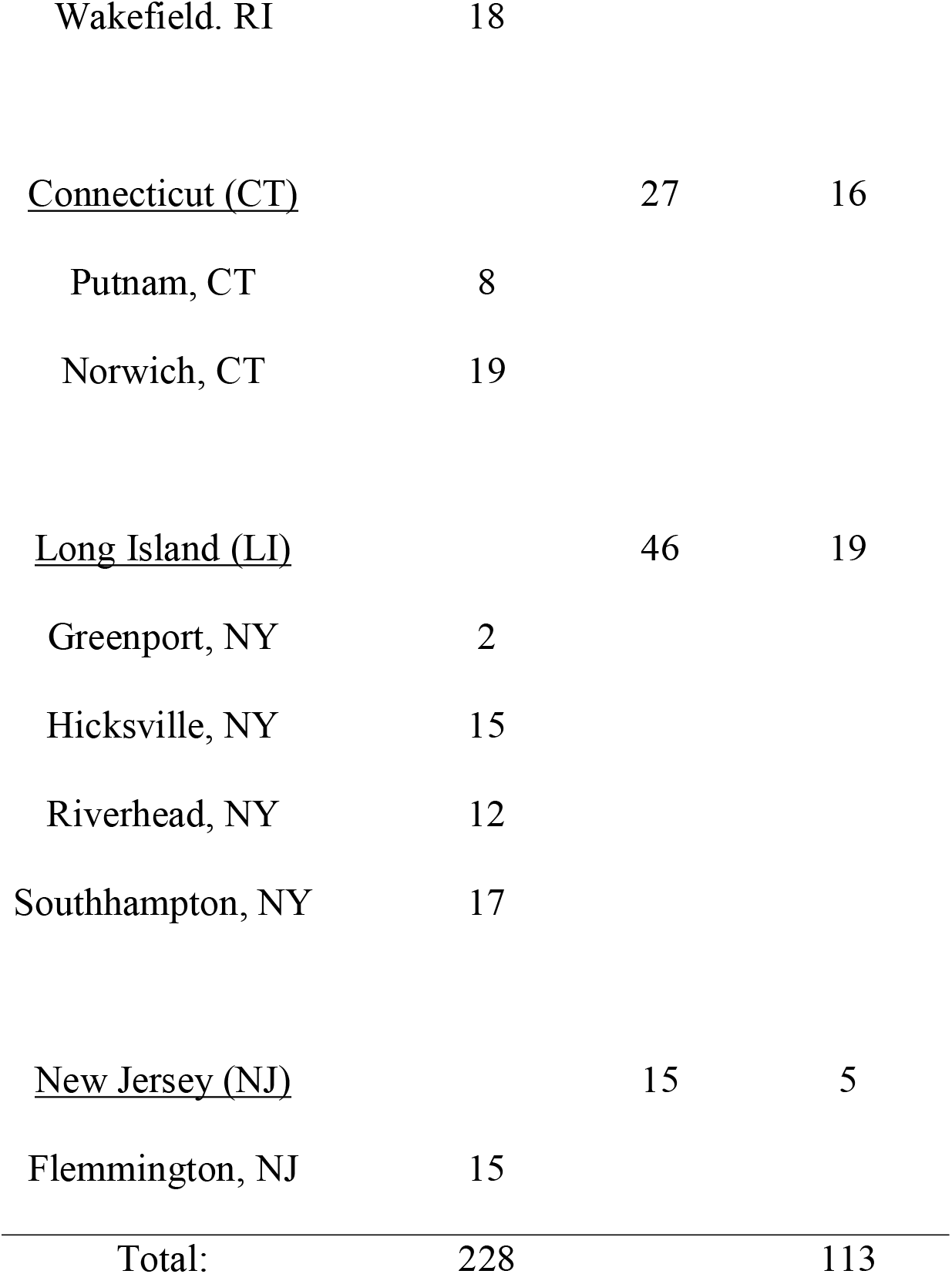
Sites from which samples were collected and the number of haplotype identified from each site.

**Table 2:** PhiPT estimates for *B. microti* from human patients by region.^a^.

|  |  | Cape | Long |  |  | New | Rhode |  |
| --- | --- | --- | --- | --- | --- | --- | --- | --- |
| Region | Boston | Cod | Island | Nantucket | Connecticut | Jersey | Island | SeMA<sup>b</sup> |
| Cape Cod | **0.51** |  |  |  |  |  |  |  |
| Long Island | 0.05 | **0.64** |  |  |  |  |  |  |
| Nantucket | **0.49** | **0.62** | **0.59** |  |  |  |  |  |
| Connecticut | 0.07 | **0.62** | 0.02 | **0.54** |  |  |  |  |
| New Jersey | 0.07 | **0.66** | 0.01 | **0.67** | 0.04 |  |  |  |
| Rhode Island | 0.03 | **0.58** | 0.04 | **0.47** | 0.01 | 0.05 |  |  |
| SeMA<sup>b</sup> | 0.04 | **0.33** | <u>0.15</u> | **0.33** | <u>0.13</u> | <u>0.14</u> | <u>0.10</u> |  |
| WMA<sup>c</sup> | 0.003 | **0.51** | <u>0.12</u> | **0.48** | 0.09 | <u>0.17</u> | 0.06 | 0.04 |

<sup>a</sup> Significant population structure $\Phi_{PT} > 0.25$ are shown in bold. Moderate population structure $\Phi_{PT} = 0.1-0.25$ is underlined.
<sup>b</sup> Southeastern Massachusetts
<sup>c</sup> Western Massachusetts

The eBurst algorithm of Phyloviz grouped the samples into 3 main clusters consisting of samples primarily from Nantucket (N population), samples primarily from Cape Cod (CC population) and those from all other sites except for SEMA (Mainland population) (Fig 3). Samples from SEMA were divided among all 3 populations. About 6% of the samples remained unresolved and were not connected to any of the 3 major groups; the majority of these (>75%) were from SEMA and RI.

**Figure 3.**
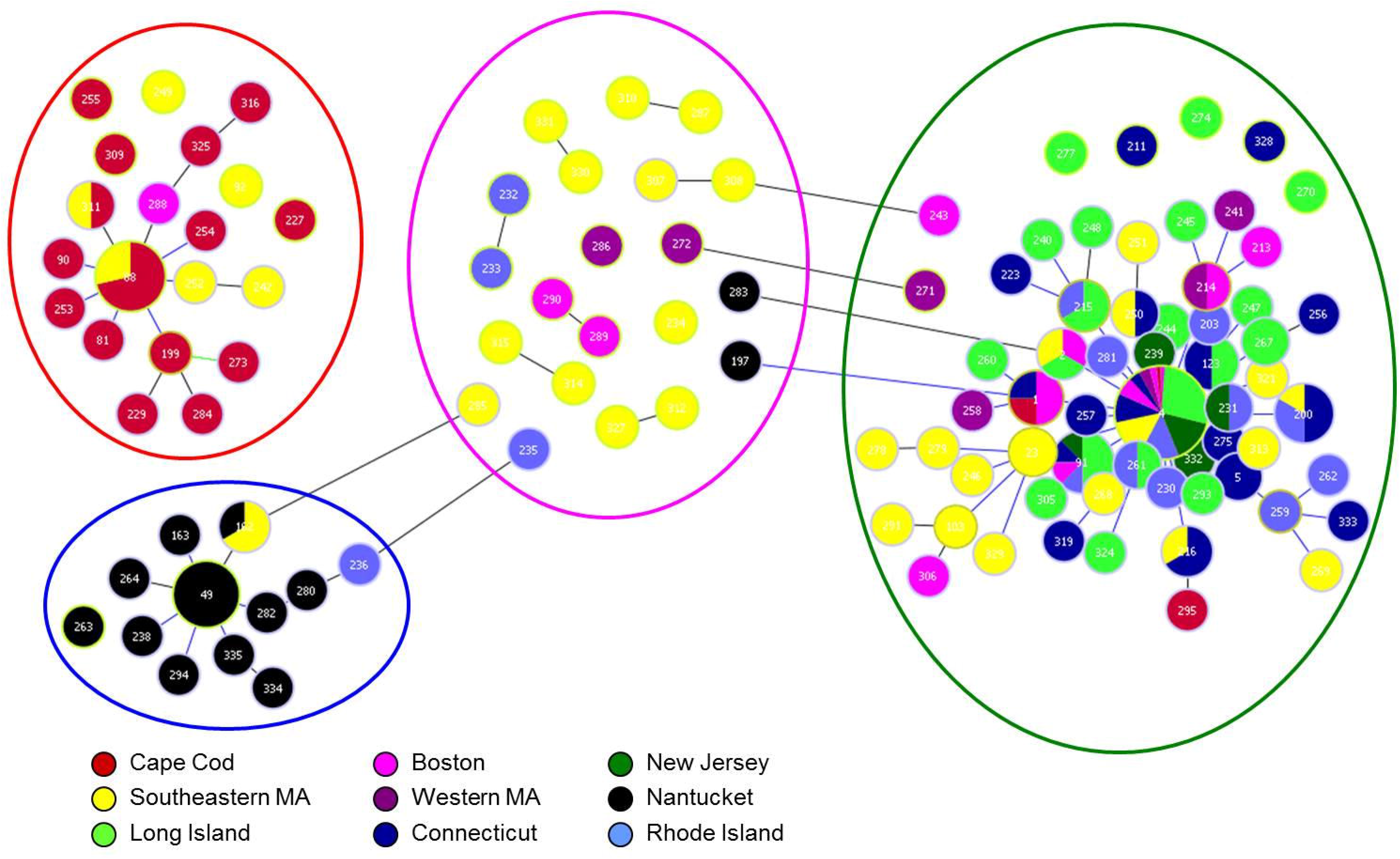
Cluster analysis of *B. microti* samples using Phyloviz. Each small bubble represents a unique haplotype. Bubbles are colored to correspond with the region from which the sample originated. The size is not directly correlated with the number of samples. Haplotypes that differ by a single locus are connected with a gray line. The large circles correspond with the population groupings calculated by TESS; blue is the Nantucket population, red is the Cape Cod population, green is mainland population and pink are the haplotypes that showed significant admixture and could not be placed solely in any of the 3 populations. Bubbles that are unconnected to the major groups are placed in the larger circles according to the ancestry coefficients from TESS.

By plotting the mean DIC for K populations from 2-8, we determined that 3 populations, K=3, best fit the data from TESS (Fig 4). Ancestry coefficients from 10 runs for K=3 were estimated for each sample, and the CLUMPP algorithm was used to combine the data from all the runs (Fig 5). These coefficients indicate the probability of membership into each of the 3 populations and corresponded well with the results from Phyloviz (Fig 3). Many samples that remained unresolved with Phyloviz showed significant amount of admixture, which would explain the inability of that algorithm to decisively place them into any single cluster (Table 3 and Fig 3 inside pink circle). However, the agreement between the two methods was not unanimous. There were a few samples that Phyloviz was unable to assign to a cluster that TESS had >85% certainty of inclusion into one of the populations (see unconnected bubbles inside larger circles Fig 3), as well as samples that Phyloviz connected to major populations that TESS could not determine to >85% probability (see bubbles with grey connections stretched to fit into pink circle Fig 3 and Table 3).

**Figure 4.**
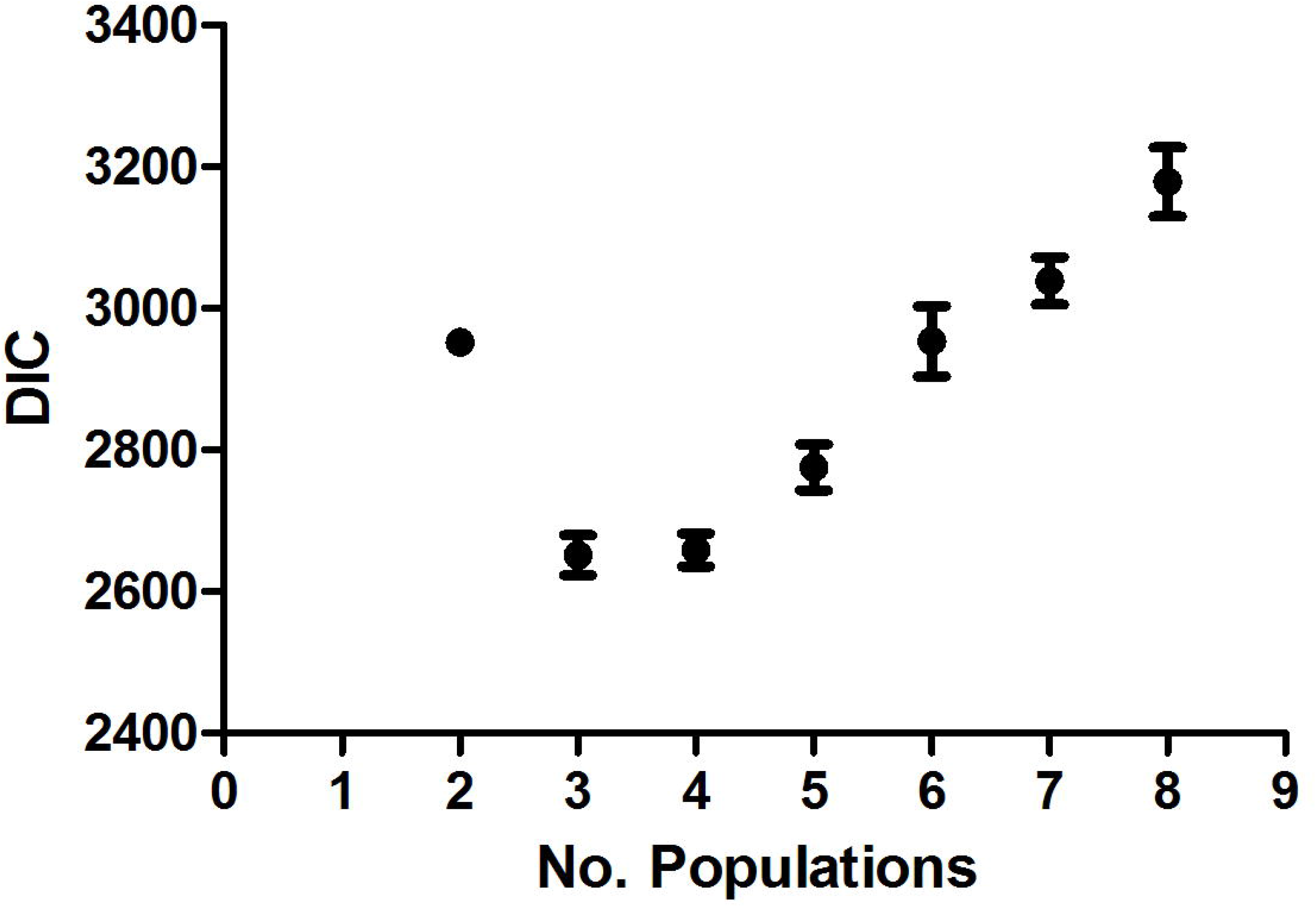
Graph of the mean DIC. Mean DIC was calculated from 10 individual TESS runs for population size 2-8. Three populations, K=3, best fit the data.

**Figure 5.**
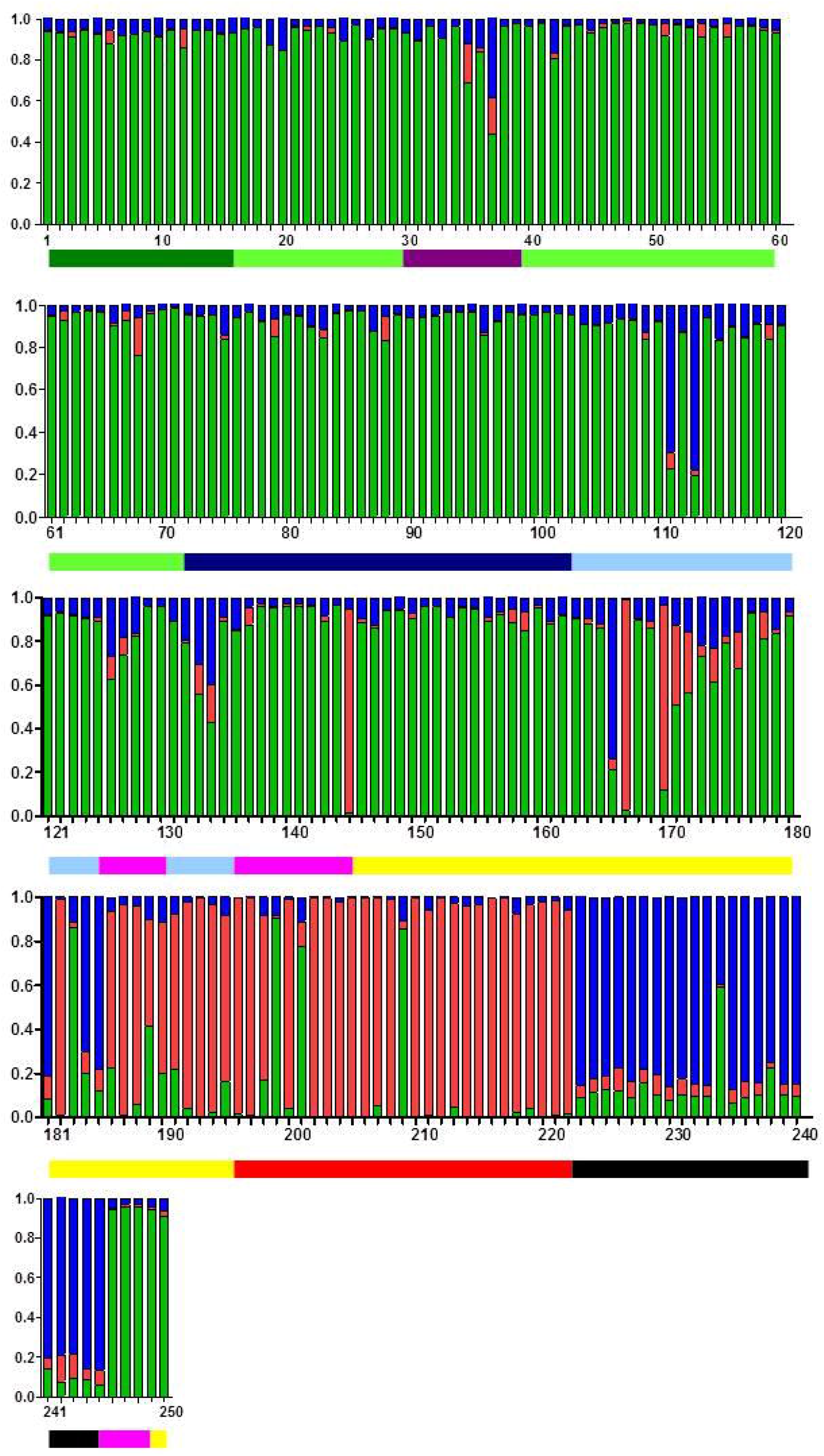
Ancestry coefficients from TESS for K=3 populations. Geographic distances between each sample point were calculated using TESS. Green corresponds to the mainland population, red is Cape Cod and blue is Nantucket. Lines beneath the bar chart indicate the source of the sample. Black= Nantucket, light blue= RI, dark blue= CT, purple= WMA, pink= Bos, red=CC, yellow= SeMA, dark green = NJ and light green= LI

**Table 3.** Ancestry coefficients from TESS of samples that showed significant admixture.

| Haplotype | Region | M | CC | N |
| --- | --- | --- | --- | --- |
| 197 | N | 0.59 | 0.01 | 0.39 |
| 272 | WMA | 0.44 | 0.18 | 0.38 |
| 286 | WMA | 0.69 | 0.19 | 0.12 |
| 315 | SEMA | 0.57 | 0.27 | 0.16 |
| 314 | SEMA | 0.51 | 0.37 | 0.13 |
| 327 | SEMA | 0.79 | 0.03 | 0.18 |
| 312 | SEMA | 0.67 | 0.17 | 0.16 |
| 307 | SEMA | 0.73 | 0.05 | 0.22 |
| 308 | SEMA | 0.62 | 0.15 | 0.23 |
| 232 | RI | 0.56 | 0.14 | 0.30 |
| 233 | RI | 0.43 | 0.17 | 0.40 |
| 289 | Bos | 0.56 | 0.14 | 0.30 |
| 290 | Bos | 0.43 | 0.17 | 0.40 |
| 331 | SEMA | 0.20 | 0.68 | 0.11 |
| 330 | SEMA | 0.41 | 0.49 | 0.10 |
| 310 | SEMA | 0.22 | 0.71 | 0.08 |
| 287 | SEMA | 0.22 | 0.71 | 0.06 |
| 235 | RI | 0.22 | 0.08 | 0.70 |
| 283 | N | 0.22 | 0.03 | 0.75 |
| 285 | SEMA | 0.20 | 0.09 | 0.70 |
| 234 | SEMA | 0.21 | 0.05 | 0.74 |

The geographically placed ancestry coefficients produced by TESS were spatially interpolated onto a map of New England (Fig 6). Haplotypes from the Nantucket population are primarily found on Nantucket. There has been some introduction into southeastern MA. The CC population also has limited scope: these haplotypes are found primarily on CC with some extending along the eastern coast of MA south of Boston. Contrary to the limited range of the N and CC populations, the mainland population dominates all of NJ, LI, CT, RI and MA, other than Cape Cod and Nantucket. It should be noted that this study did not include any data from Martha’s Vineyard; so it may be that the predicted populations included in this figure are erroneous.

**Figure 6.**
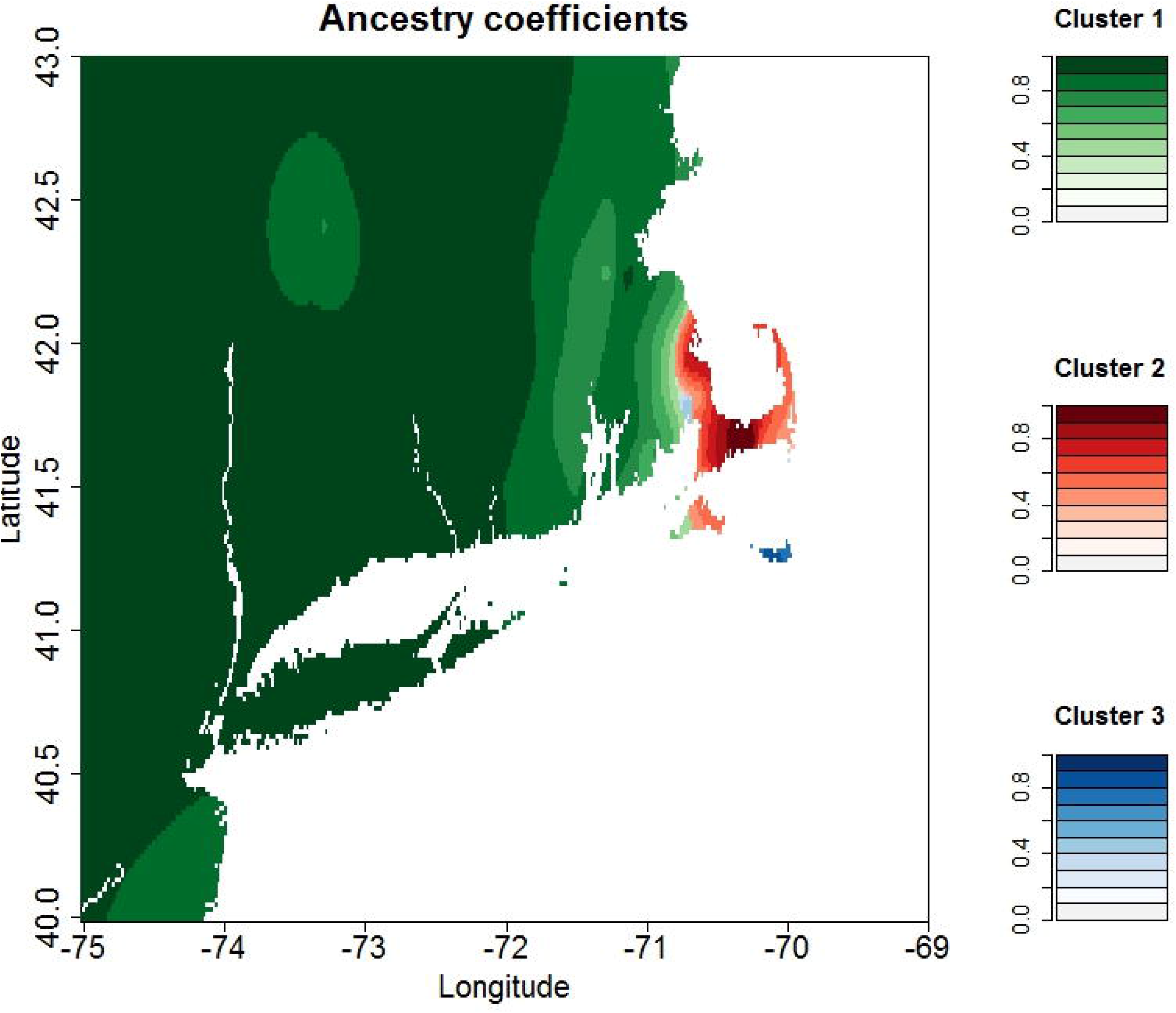
Geographic interpolation of the ancestry coefficients showing the distribution of each population of *B. microti.* Cluster 1(green) = mainland population, cluster 2 (red) = Cape Cod population, and cluster 3 (Blue) = Nantucket population. Areas with samples that have a high admixture coefficient, ie a high probability of membership to that population, are shaded darker. Lighter shades indicate areas where there the ancestry coefficients are lower, indicating areas where mixing is occurring. This study did not include data from Martha’s Vineyard; so the predicted populations on that island may be erroneous.

Each of the 3 populations has a dominant haplotype that is also the putative ancestral type (as determined by Phyloviz), type 4 for mainland, type 49 for Nantucket, and type 88 for Cape Cod (Table 4). Type 49 is present in 48% of Nantucket samples; Type 88 is found in 37% of Cape Cod samples, and type 4 ranges from 33% to 75% in the regions included in the mainland population (Figure 7). SEMA is the only region with a mixture of the dominant types; type 4 was detected in 22% of samples and type 88 detected in 7%. All other haplotypes in this study are detected only once or twice from any given region, with the exception of type 91 from LI which was found 4 times (8% of the observed haplotypes). Type 91 differs from the dominant type 4 by only the BMV1 locus (335bp instead of 340bp) of type 4.

**Figure 7.**
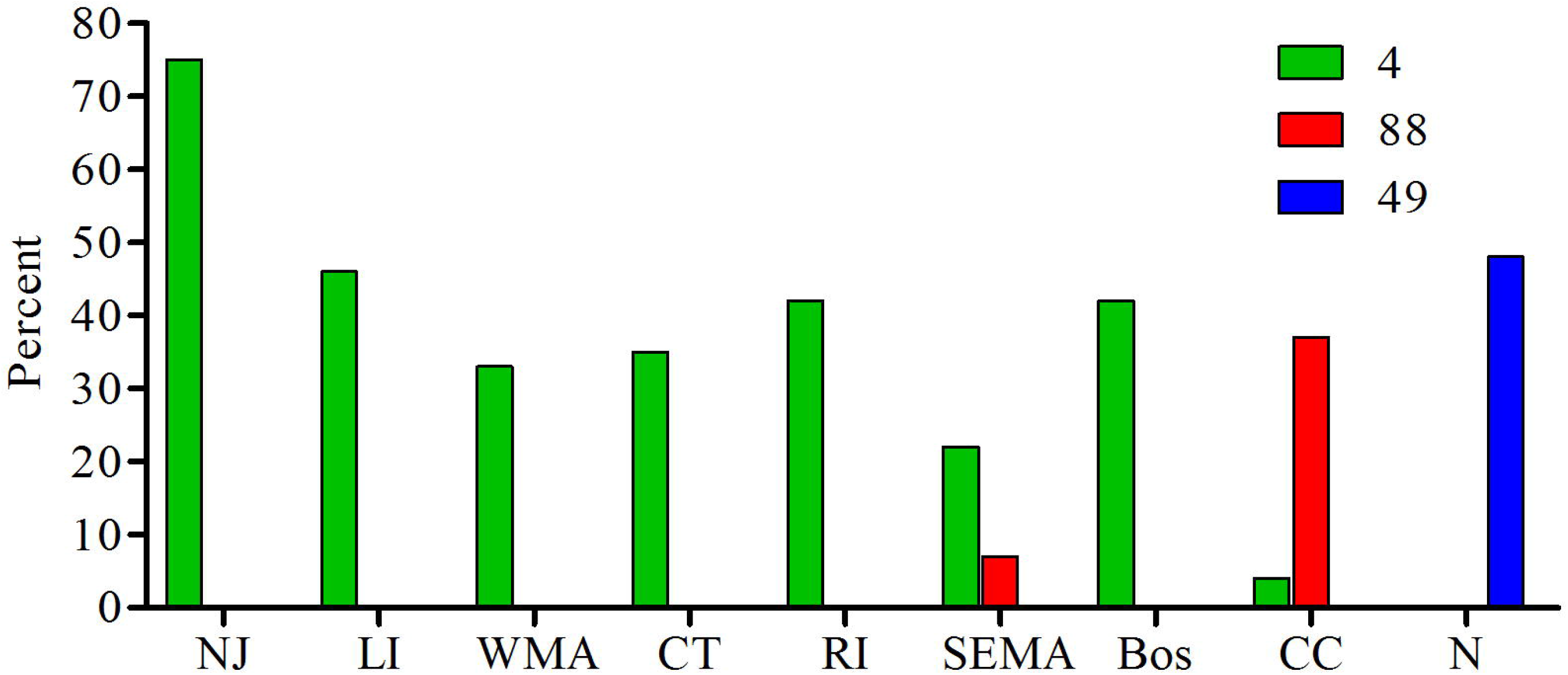
The percent of the total samples for each region for each of the main haplotypes: type 4, type 88 and type 49.

**Table 4.**
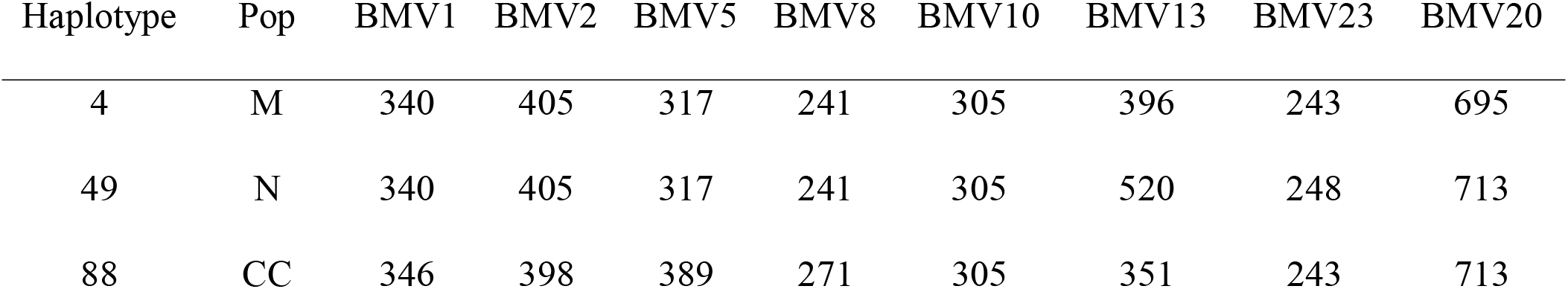
The microsatellite amplicon sizes of the 3 major haplotypes in base pairs.

## Discussion

Our analysis provides data to help reconstruct the tempo and mode of the processes that have led to the current epidemic population structure of *B. microti* in northeastern US. There are at least 3 distinct populations of *B. microti* in New England, as we suggested previously [27, 28] in analyses of ecological as well as clinical samples, with PhiPT ranging from 0.32-0.67 between them (Table 2). Each of the three populations has a single dominant haplotype that is found in at least 30% of the samples from each site and is the presumed ancestral strain; type 4 for mainland, type 49 for Nantucket and type 88 for CC. Southeastern MA is currently experiencing a natural experiment as the 3 populations, CC, N and M, are zoonotic in this area. The CC population is moving northward and westward along the eastern coast of MA, the N type is invading from the southern coast, and the mainland type is invading from the west. The genetic signature from all 3 populations can be clearly detected in clinical samples from this area, and significant admixture is occurring (Fig 6). For this reason, the diversity of *B. microti* from SEMA is significantly greater than that from all other regions in our study. Although we do not know when each of the *B. microti* populations were first introduced into SEMA, nor which one arrived first, type 4 is found more often in this area and the majority of samples harbor loci that originate from type 4. This dominance is clearly represented in the map of the ancestry coefficients from TESS, and suggests that type 4 parasites have some attribute that allows for greater amplification than do the other *B. microti* populations. It may be that type 4 parasites are more transmissible.

If the expansion of *B. microti* in New England was caused by individual founders “pulling” the population, we would have expected the diversity estimates from ancestral sites (Nantucket; Cape Cod; Long Island; [11], where cases have been diagnosed since the 1970s, to be greater than those from incipient sites with more recent emergence of cases. However, this was not the case; the diversity estimates of *B. microti* from the regions we sampled across the northeastern United States were not significantly different. In fact, the diversity of *B. microti* from ancestral sites, such as Nantucket and Long Island, were no greater than those from more newly established sites. Furthermore, the diversity from coastal CT was not significantly different than that from northern CT where babesiosis cases were first detected 15 years later. The maintenance of diversity across New England supports the theory that expansion was the result of a “pushing” population expansion, consistent with the stepping-stone hypothesis inferred by Walter and colleagues [8]. Notably different, however, were samples from NJ; their diversity was significantly less than those from every other site in our study; more than 70% of the parasite samples comprised the dominant type 4. The lack of genetic diversity is consistent with the New Jersey foci representing newly established populations that have experienced significant founder effects. However, *B.* microti-infected ticks were documented from northern New Jersey in the early 1990s [38] and human cases shortly thereafter [39]. New Jersey became endemic for babesiosis at the same time as northern CT and northern RI, but the diversity of *B. microti* from those states are similar to those from the rest of the study populations. The biological basis for the limited diversity found in New Jersey *B. microti* samples remains to be described.

Some patient samples may have been mistakenly assigned to location because we used convenience samples that were de-identified other than for site of the contributing clinical practice. We assumed that a case became exposed near the healthcare provider who provided the sample to Imugen for analysis. Residents of any of our sites are likely to travel within the northeast, and may vacation or visit in sites where risk is similar to where they live. We are confident, for example, that two samples from our Nantucket cohort acquired infection elsewhere. Each of these samples contained parasite haplotypes that grouped with the mainland population. We have analyzed sufficient numbers of ecological samples from Nantucket Island and have never detected the other lineages [27]. Despite this clear example of mistaken assignment, the outcome of our analysis did not appear to be effected; TESS correctly concluded that Nantucket Island is dominated solely by the Nantucket population and the other sites by their respective parasite populations. Accordingly, we believe that our analysis is robust enough to be unaffected by other unknown errors in geographic assignment of samples and that our conclusions about the population structure of *B. microti* in the northeastern U.S. are reasonable.

It is also possible that focusing our analysis solely on parasites derived from presumably symptomatic patients (those presenting to a healthcare provider who in turn requested analysis of a sample for confirmation of a diagnosis) does not capture variation of all those that may be present in the enzootic cycle of the mainland parasites. There is as yet no published evidence that the diversity of *B. microti* infectious for humans differs from that in local mice or ticks, i.e., that only a subset of naturally occurring strains are zoonotic. However, such an argument would need to apply across all sites and we note that there is much variation evident in parasites from patients presenting to healthcare providers on Nantucket, Cape Cod, or Southeastern Massachusetts.

Significant differentiation (PhiPT >0.36) between each of the 3 populations implies that they have been isolated from each other and remain so. We have previously speculated that the microbial guild transmitted by *I. dammini* had been maintained in relict or refugial foci during glaciation [11]. Then too, postcolonial deforestation likely provided a fragmented landscape that only allowed for perpetuation of ticks and their hosts in small less-disturbed natural foci. The lack of differentiation among parasites from the mainland sites, from central NJ westward to RI, appears to be inconsistent with a scenario of multiple relict foci across the mainland northeastern landscape, with coalescence of the isolated demes occurring as a result of amplification and expansion of the foci as successional habitat increased over the last 100 years. In the 1990s, babesiosis was documented from 3 distinct sites within the area where the mainland population parasites have been detected, viz., Long Island, southeastern CT and central NJ. Each of these foci was isolated from the others; few cases were identified in areas between them. Ecological sampling, where it was done, supports the inference that *B. microti* was indeed absent or very rare [7,13,15,16,18,40]. We expected to detect a distinct genetic signature of multiple small isolated foci within parasites from the mainland lineages but there is little differentiation among LI, CT, RI and NJ, and our analyses group these sites together into a single population. In fact, the mainland haplotype, type 4, dominates from NJ eastward through NY, CT and RI and northward towards Boston, creating an epidemic population structure.

It may be that these sites were not isolated for sufficient time for genetic drift to operate, thereby explaining the lack of differentiation among mainland parasites. It is also possible that the epidemic population structure occurred purely by chance, i.e. genetic drift has occurred as *B. microti* has expanded leading to an overabundance of a single haplotype. Some alleles may reach a high frequency because of repeated founder events [22], a process called genetic surfing [26]. We assume that our VNTR loci are neutral or are not linked with loci under selection and thus the observed lack of variation is not due to selective constraints. The alternative hypothesis for the lack of diversity among mainland *B. microti* is that there were no refugial or relictual sites within fragments of forest, and that the parasite populations have not actually been isolated from each other, allowing sufficient gene flow within the various sites comprising the mainland. However, the population structure of *I. dammini* suggests otherwise. A seminal study of the population structure of this vector tick and *B. burgdorferi* infecting them [41] sampled 12 sites in the northeast from Massachusetts to Virginia; 5 of these overlap with our area of study. Mitochondrial 16SrDNA haplotypes demonstrated that the New York-CT region may have contained refugial tick populations that served as a source for expansion of the range of *I. dammini.* Although tick populations that were sampled were structured, this was not observed for *B. burgdorferi,* although the borrelial genes that were analyzed were likely to have been influenced by balancing selection [41]. Additional studies are required to identify the relative contributions of selective and demographic processes that serve as the basis for biogeographic variation in northeastern populations of *B. microti*.

We believe the most likely scenario is that type 4 parasites have selectively swept across the mainland landscape, replacing and erasing historic genetic signatures of other lineages. Such a hypothesis is not without precedent with the microbial guild maintained by *I.* persulcatus-like ticks. The population structure of *B. afzelii* (an Eurasian agent of Lyme disease that appears restricted to rodent hosts) in Sweden is essentially clonal, which may be the result of the epidemic spread of a single genotype [42]. Across Europe, however, *B. afzelii* has significant population structure [43], similar to what we have found in this study. There are likely public health implications of a specific *B. microti* lineage that appears to be rapidly expanding its range.

## Acknowledgements

Many clinicians and clinical practices submit diagnostic samples to Imugen Inc for testing. Samples for this study were de-identified and thus we do not know the identities of their submitters, but we thank them for their contribution to this study.

